# Sex-specific immunocompetence: resistance and tolerance can both be futile but not under the same circumstances

**DOI:** 10.1101/2024.06.10.598263

**Authors:** Franziska A. Brenninger, Viktor Kovalov, Hanna Kokko

**Author notes:** Shared first coauthor (Franziska A. Brenninger, Viktor Kovalov). **Author contributions** Conceptualization – all the authors; building and investigating the model – all the authors; writing original draft – F.A.B. and V.K.; reviewing and editing – all the authors; supervision – H.K. All the authors read the final version of the manuscript and approved it for the publication. **Funding** F.A.B. and V.K. were funded by University of Zurich, H.K. acknowledges funding by the Alexander von Humboldt Foundation. **Data & Code availability** MATLAB files with the model code and scripts used to generate data and figures will be freely available after the manuscript is accepted for publication. The generated data are visually represented in figures in the main text and in the supplementary materials. For review purposes, all the MATLAB files can be requested with the following link: https://zenodo.org/records/11548626.

## Abstract

Immunocompetence evolution can involve a ‘resistance is futile’ scenario, if parasite encounter rates are so high that high investment in resistance only marginally delays infection. Here, we investigate two understudied aspects of ‘futility’. First, immunocompetence is usefully categorized as reducing the rate of becoming infected (resistance) or reducing the negative fitness consequences of infection once it happened (tolerance). We compare the prospects of futility for resistance, tolerance, and their joint occurrence, showing that resistance futility arises with respect to parasite encounter rates, while tolerance futility arises with respect to parasite virulence. However, if the same host trait improves pleiotropically both resistance and tolerance, futility disappears altogether and immunity investment remains profitable when increasing parasite encounter rates, virulence, or both. Second, we examine how sexual selection strength impacts these findings. If one sex (typically males) is near the faster end of a fast-slow continuum of life histories, then life history patterns reflecting futility can evolve sex-specificity. The solutions often feature sexual dimorphism in immunocompetence, but not always in the direction of strong sexual selection yielding low immunity: sexual selection can select for faster and ‘sicker’ lives, but if sexual selection also causes traits that impact parasite encounter rates, the results are strongly dependent on whether futility (along any axis) plays a role.

**Lay Summary:** Intuition suggests that investment into immunity is higher, when hosts frequently encounter parasites. While there are examples that confirm this, in other cases, hosts have been shown to abandon immune defenses under high parasite pressure. We reconcile these findings by modelling the optimal host resource allocation towards immunity under varying parasite pressure and strength of sexual selection. Our results show two axes along which immunity investments are futile and should therefore be abandoned in favor of investing into reproduction: resisting infection becomes futile under high parasite abundance, while tolerating the harmful effects of infection is not beneficial under ever increasing parasitic virulence. However, investments of organisms that are capable of both resistance and tolerance mechanisms yield fitness payoffs also when parasites are highly virulent and abundant. This work highlights the impact of parasites and immune defenses on optimal immunity investment levels in hosts, an insight which also complements theory on sex-specific immunity.

## Introduction

Hosts employ various strategies to fend off parasites, ranging from behavioural avoidance (Gibson & Amoroso, 2022), to a variety of physiological mechanisms summarizable as the immune system (Nicholson, 2016). Immunocompetence, i.e., the ability of a host to resist the establishment of a parasite and/or to limit and possibly eradicate parasite growth once infected, can be broadly divided into resistance and tolerance. Resistance refers to any immune mechanism reducing the infection probability after encountering a parasite, while tolerance captures the host’s ability to minimise parasite-imposed harm once infected (Råberg *et al*., 2009, Martins *et al*., 2019). Resistance and tolerance create distinct host-parasite dynamics (Schneider & Ayres, 2008; Vincent & Sharp, 2014; Miller *et al*., 2007; Klemme *et al*., 2022), though a host may also combine both: resisting infection as long as it can, employing tolerance afterwards (Råberg *et al*., 2009). Note, however, that terminology varies among authors: Miller *et al*. (2007) define tolerance as above but consider it a subcategory of resistance, Jokela *et al*. (2000) use tolerance to refer to a scenario where any type of defence has evolved to be low, and Best *et al*. (2010) use resistance and tolerance to refer to two different traits that both take effect in infected hosts.

An interesting theoretical finding is a scenario where “resistance is futile”. Futility is typically discussed with respect to resistance: if parasite encounter rates are so high that strongly resistant hosts still become infected, then, assuming a trade-off between resistance and current reproduction, reduced investment into resistance is selected for (Jokela *et al*. 2000; Walsman *et al*., 2023). This may involve ‘classical’ resistance, as well as any behavioural avoidance of parasites (Walsman *et al*., 2022, 2023). The “futility” cases are a counterexample to the intuition that more abundant parasites lead to stronger host immune defences (Lindström *et al*., 2004).

Here, we investigate two understudied aspects of ‘futility’. First, futility usually involves evolutionary scenarios where high parasite encounter rates select for little or no resistance. The parallels to the tolerance evolution are scarcely discussed, though it is known that tolerance can be lost under certain conditions, such as short host lifespans (Miller *et al*., 2007). We aim to compare the prospects of futility for resistance, tolerance, and their joint occurrence.

Second, we examine how sexual selection strength impacts these findings. The link to the concept of futility is clear: by “giving up” resistance when it is futile, a host makes the most of its (potentially brief) time being still parasite-free. If we assume that males optimise life in “live fast, die young” manner (differing from females in their placement on “fast-vs-slow” life-history axis (Kokko 2024), a “resistance is futile” response can show sex-specificity.

Sexually dimorphic immunocompetence (Zuk *et al*., 2004) has been reported in birds (Møller *et al*., 1998; Vincze *et al*., 2022, Sauer *et al*., 2024), mammals (Moore & Wilson, 2002; Klein, 2012; Wilkinson *et al*., 2022; Guerra-Silveira & Abad-Franch, 2013), fishes (Shepherd *et al*., 2012; Dong *et al*., 2017) and insects (Kurtz *et al*., 2000; Belmonte *et al*., 2020); for a review see Kelly *et al*. (2018). Although males are not always the ‘sicker sex’ (Sheridan *et al*., 2000; Stoehr & Kokko 2006; Hillegas *et al*., 2008; Sanchez *et al*., 2011), there is a general expectation that strong sexual selection should make males favour high mating success even if it comes with a shortened lifespan. The links to futility, however, remain unexplored.

We present a model that assumes (1) a trade-off between immunity and reproduction, (2) varying levels of sexual selection strength, (3) variation in parasite encounter rate and virulence, and (4) two immune mechanisms – resistance and tolerance, whose impact is assessed separately as well as jointly. We consider the effects of resistance and tolerance on the immunity-mediated trade-off between reproduction and lifespan, together with the effects of sexual selection.

### Model

Our continuous-time model optimizes the allocation of resources *x* (0 ≤ *x* ≤ 1) into immunity that trades off with reproduction. Resources not allocated towards immunity (1–*x*) are invested into reproduction. A host starts its life in a susceptible state (S) with baseline mortality *μ* and parasite encounter rate *k* > 0. Every parasite encounter involves a chance of transitioning to the infected state (I). Parasite virulence *ω* > 0 elevates the mortality of infected hosts.

We consider three scenarios. In the *resistance* scenario, investment into immunity *x* reduces infection probability upon parasite encounter. In the *tolerance* scenario, *x* instead mitigates harm imposed by parasites after infection. Finally, in the *pleiotropic* scenario, *x* feeds into both immunity mechanisms simultaneously.

For each scenario, we denote the expected lifespan *L* of a host capable of resistance (R), tolerance (T) or both (P, for pleiotropy) as the sum of the time a host is expected to spend in the S and the I states:

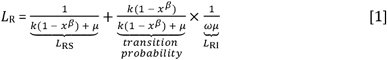

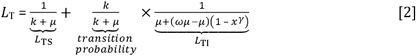

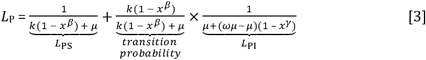

In scenarios involving resistance (equations 1 & 3), the probability of infection upon each parasite encounter equals 1–*x*^*β*^, *whe*re lower values of *β* imply more efficient reduction in risk (Figure S1b). If *β* is high, *s*ubstantial allocation (*x*) is required to yield a considerable reduction. These assumptions translate into a transition rate *k*(1–*x*^*β*^) from S to I.

Remaining in the S state requires avoiding both infection and death. In cases where immunocompetence *x* improves resistance, the expected time spent in S is longer if *x* is high (equations 1 & 3), while if *x* only confers tolerance (equation 2), the time to infection depends on *k* without involving *x*. In all settings, the expected time spent in a S state is also limited by death rates *μ*.

The expected time spent in the I state depends on lifetime probability of transitioning from S state to I state in the first place (second term in equations 1-3), multiplied by the expected lifespan conditional on ever entering that state (the last terms in equations 1-3). After infection (but before dying), a host remains some time in the I state. Since we do not include recovery in our model, the only way to leave the I state is through death. In scenarios with no tolerance, virulence *ω* elevates the death rate from *μ* to *ωμ* and thus the expected lifespan from the point of infection onwards is 1/(*ωμ*). In scenarios including tolerance (equations 2 & 3), immunity investment *x* mitigates the mortality changes caused by virulence: we assume it diminishes the difference between mortality rates while infected and uninfected (*ωμ* – *μ* in the absence of tolerance) through a multiplication by 1–*x*^*γ*^, where *γ* modulates how allocation *x* translates into longer lifespan once infected (*γ* >0) (Figure S1c). Thus, mortality changes from *μ* to *μ+*(1–*x*^*γ*^)(*ωμ*–*μ*) upon infection. Note that even with mortality reduction through tolerance, mortality in the I state does not fall below mortality in the S state.

We assume that host reproductive success is contingent on the fraction of resources invested into reproduction (1–*x*), both in the S and in the I states. Reproductive output per time unit equals (1–*x*)^*α*^, *whe*re *α* describes the strength of sexual selection (*α*>*0)*. High values of *α* indicates strong sexual selection, requiring a large investment into reproductive effort before reproductive success increases appreciably (Figure S1a). This reflects a scenario where strongest investment into reproduction yields access to many matings, while competitors with less investment fail to reproduce. Low values of *α*, conversely, imply that relatively small investment into reproduction suffice to yield substantial reproductive success, and increasing investment further yields diminishing returns — as is often the case for females.

The expected fitness of a host of a specific sex is a product of reproductive output per time unit multiplied with the expected lifespan. For each scenario, ‘resistance’, ‘tolerance’ and ‘pleiotropic’, this yields the following equations describing the expected lifetime fitness:

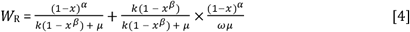

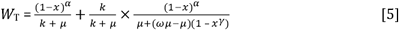

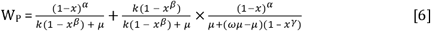

We numerically derive the optimal immunity investment *x* that maximizes fitness for a host, and then interpret the resulting dependencies on *α, β* and *γ*.

## Results

### The effect of parasite encounter rate on optimal immunocompetence depends on the immunity mechanism

When reproduction trades off with resistance only, lowest and highest parasite encounter rates *k* produce similar outcomes, where investment into immunity remains low (red dots near the left end of the *x* axis, Figure 1). It is easy to understand why immunity is neglected with low parasite encounter rates – most lives end without entering the infected state, thus, no incentives to invest into immunity. However, immunity remains low also at high parasite encounter rate with lifespans evolving to be short. This is a classic example of a “resistance is futile” strategy: when parasitic encounters are so frequent that even high immunity cannot prevent infection for any substantial amount of time, it becomes optimal to neglect immunity and maximise offspring production before death. Substantial immunity investment in the resistance scenario only evolves at intermediate values of parasite encounter rate *k* (Figure 1).

**Figure 1:**
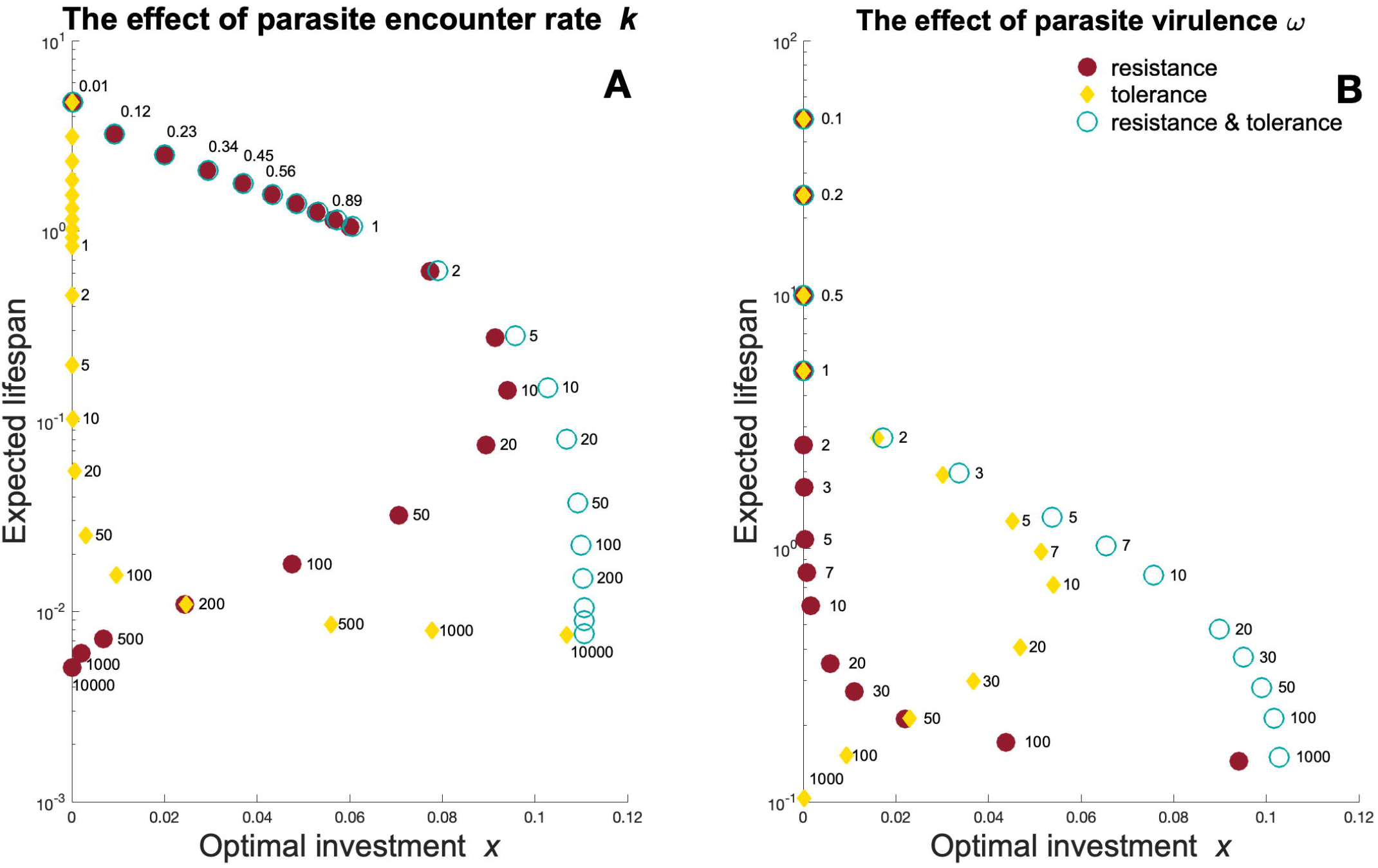
Optimal investment *x* and its consequence, the expected lifespan (sum of expected lifespan in two states, susceptible and infected), as a function of (A) parasite encounter rate *k* and (B) parasite virulence *ω*, with values of *k* (for A) and *ω* (for B) indicated as numbers next to each dot. Dot colour and shape indicate immunity mechanisms included into a given model scenario: red dots – ‘resistance’ scenario, yellow diamonds – ‘tolerance’ scenario, blue circles – ‘pleiotropy scenario where immunocompetence improves both resistance and tolerance. Other model parameters: *α* = 2; *β* = 0.5; *γ* = 0.5; *μ* = 0.2; in (A), *k* = 10^−2^…10^4^ and *ω* = 1000; in (B), *k* = 10 and *ω* = 10^−1^…10^3^.

With tolerance, benefits from immunity investment are based on decreasing the additional mortality induced by parasites. There is no “resistance is futile” pattern for tolerance (Figure 1) with respect to parasite encounter rates; instead, allocation into tolerance-conferring immunity increases with parasite encounter rate. Note that this increase only mitigates and does not prevent the lifespan loss associated with parasitism. Taken together, this yields a pattern where the highest immunity investment associates with lowest lifespans in the tolerance scenario (yellow squares, Figure 1).

When the same investment improves both resistance and tolerance (pleiotropy), optimal immunity investment is high under intermediate and high parasite encounter rates (blue circles, Figure 1). The highest investment again associates with the lowest lifespan, resembling the tolerance scenario but deviating from the resistance scenario, where the highest allocation was found at intermediate lifespans. With highest parasite encounter rates, optimal immunity investment does not plummet, as tolerance mitigates any additional infection-related mortality caused by infection (Figure 1).

### Futility in response to parasite encounter rate, virulence, or both?

The above example shows interesting patterns, but was drawn for one specific value of virulence *ω* (as well as baseline mortality *μ*, as listed in Figure 1). Broadening the view to other values of *ω* is necessary to infer generality – while *μ* merely scales the lifespan outcomes and its precise value is less important.

Exploring solutions across varying parasite encounter rate *k* and virulence *ω* allows asking whether either parameter promotes ‘futility’. Along the encounter rate axis, allocation diminishes (Figure 2A: colours turn darker) when moving in the direction of increasing *k*, but only for the resistance scenario; here it operates irrespective of virulence, *ω*.

**Figure 2:**
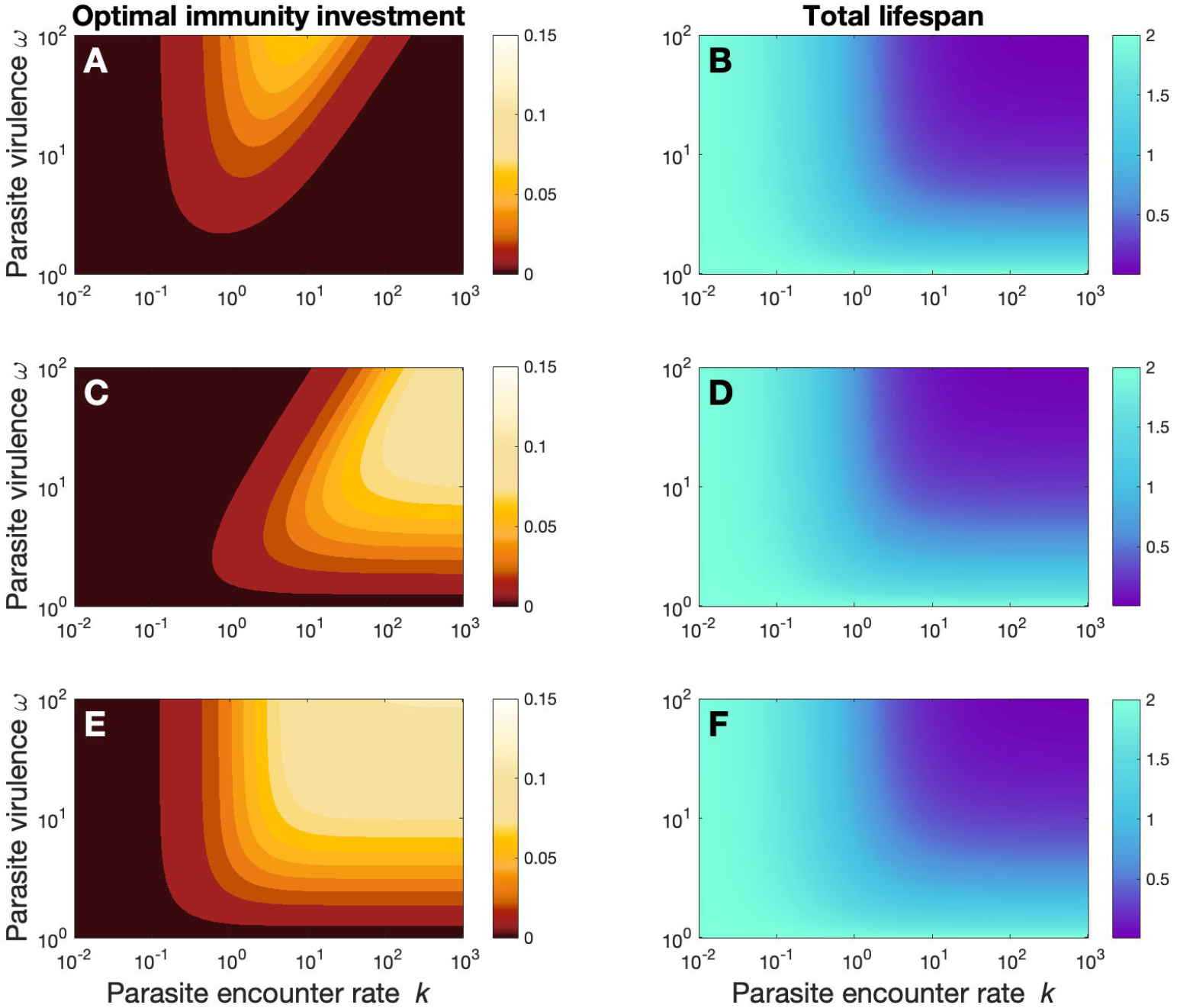
The effect of parasite virulence and parasite encounter rate on (A, C, E) optimal immunity investment and (B, D, F) expected lifespan. Immunocompetence is assumed to improve resistance (A, B), tolerance (C, D), or both (E, F). The investment heatmaps (A, C, E) show the optimal immunity investment *x* for each combination of parasite encounter rate *k* (x-axis, log scaled) and parasite virulence *ω* (y-axis), with *x* represented as a gradient from dark red (*x* = 0) to white (*x* = 0.15). The lifespan heatmaps (B, D, F) assume individuals invest according to the optimal value *x*, depicting expected lifespan as a gradient from dark violet (*L* ≈ 0) to cyan (*L* = 2). Parameter values: *α* = 2; *β* = 0.5; *γ* = 0.5; *k* = 10^−2^…10^3^; *μ* = 0.5; *ω* = 10^0^…10^2^.

The effect of virulence *ω* on optimal investment strategies differs markedly between scenarios (Figure 2A,C,E). In the resistance scenario, increasing virulence heightens allocation into immunity (Figure 2A, moving from low to high values along the y-axis makes colours lighter) with an exception for very low parasite encounter rate. In this case, allocation remains low regardless of parasite virulence. Neither the common cases, nor the exception involving low encounter rates, show futility.

The pattern is the exact opposite for tolerance: immunity investment decreases with increasing virulence (colours darken when moving from bottom to top, Figure 2C), indicating futility along the vertical axis (Figure 2C) when immunocompetence improves tolerance. While resistance futility required highly abundant parasites, tolerance futility occurs with highly virulent parasites. If parasites significantly shorten life, and significant mitigation via tolerance is unachievable, the optimal strategy maximizes fitness with a short life and fast reproduction, rather than prolonging it through tolerance.

Finally, when immunity investment improves both tolerance and resistance, the question is whether the system combines both types of ‘futility’, only one, or none. The answer is ‘none’ (Figure 2E), but with the subtlety that there are strong responses of increasing investment when the parasite encounter rate increases, while the responses to virulence only change when moving from low virulence to somewhat higher values and do not change much thereafter.

Like Figure 1, different immunity scenarios yield similar results for the expected host lifespan (Figure 2B,D,F). Total lifespan diminishes with increasing parasite encounter rate as well as with virulence (dark blue areas at the top-right side of Figure 2B,D,F), despite adaptive changes in immunocompetence that differ between resistance, tolerance, and pleiotropic scenarios (Figure 2A,C,E).

### Stronger sexual selection leads to weaker immunity, but the net effect depends on parasite encounter rates

Sexual selection strength interplays with parasite abundance to influence optimal immunocompetence (Figure 3). In general, the relationship sexual selection strength and investment into immunity is negative (Figure 3: colours darken when sexual selection increases). This is logical: relaxed sexual selection (*α* < 1) implies diminishing returns of reproductive success with ever-higher investment in reproductive traits, rendering the lifespan-extending trait of immunocompetence relatively more beneficial. However, at very low parasite encounter rates, immunity investment remains low regardless of sexual selection strength, as infection risk remains minimal.

**Figure 3:**
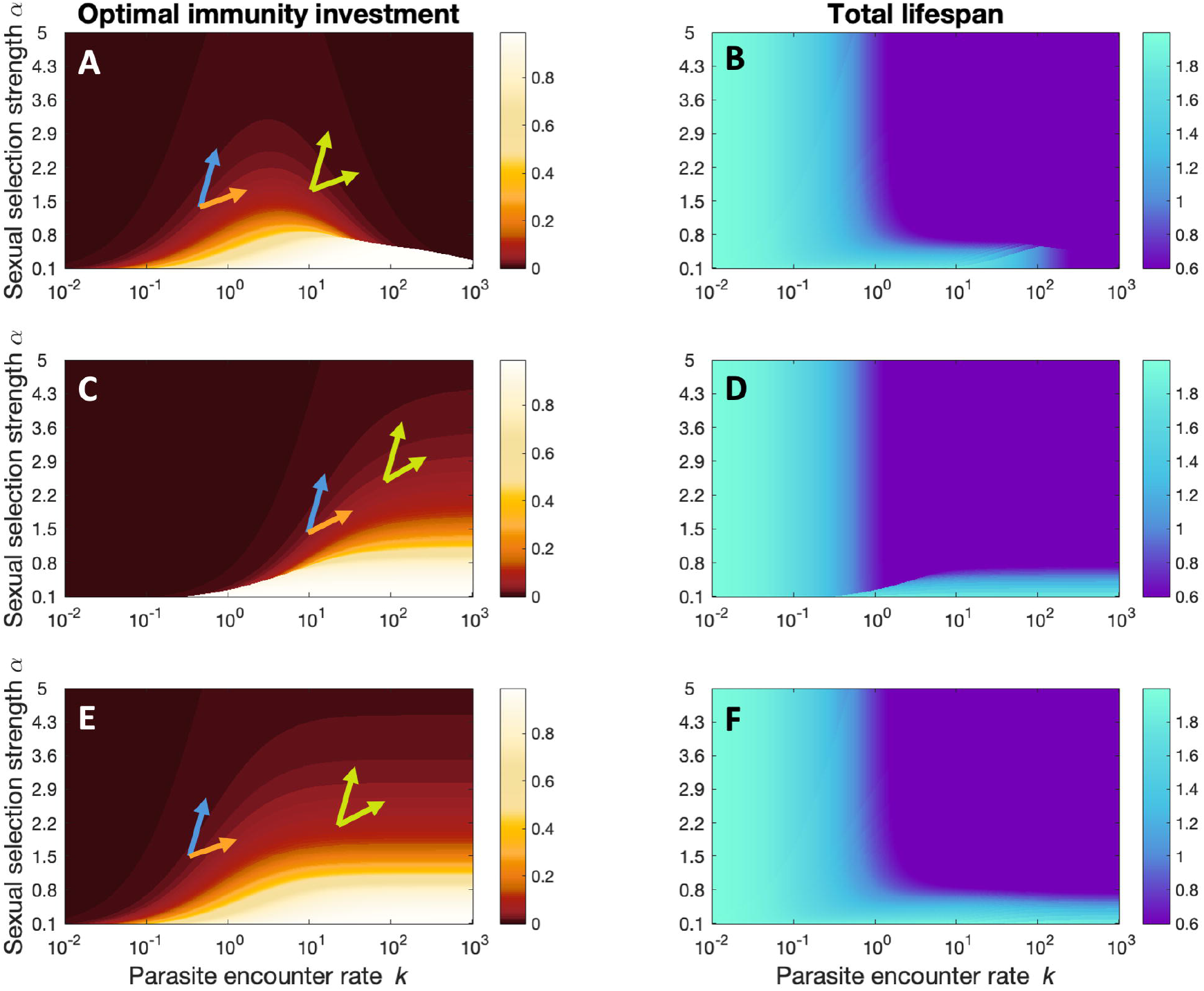
The effect of strength of sexual selection and parasite encounter rate on optimal immunity investment (A, C, E) and expected lifespan (B, D, F). Immunocompetence is assumed to improve resistance (A, B), tolerance (C, D), or both (E, F). The investment heatmaps (A, C, E) show the optimal immunity investment *x* for each combination of parasite encounter rate *k* (x-axis, log scaled) and strength of sexual selection *α* (y-axis), with *x* represented as a gradient from dark red (*x* = 0) to white (*x* ∼ 0.99, the maximum value in this model setup). The lifespan heatmaps (B, D, F) assume individuals invest according to the optimal value *x*, depicting expected lifespan as a gradient from dark violet (*L* = 0.6) to cyan (*L* = 2). Arrows inside the heatmaps point out locations referred to in the main text. All arrows point towards areas where both *α* and *k* increase compared to the arrow’s point of origin. Only yellow arrow points at higher optimal immunity investment. Model parameters: *α* = 0.1…5; *β* = 0.5; *γ* = 0.5; *k* = 10^−2^…10^3^; *μ* = 0.5; *ω* = 30.

We do not explicitly specify ‘males’ and ‘females’ in our model. Instead, we consider their situation to differ in two ways: the sex experiencing stronger sexual selection (often males) is located higher up along the *α* axis (Figure 3); additionally, some sexually selected traits, e.g. fighting and increased mobility, may elevate the parasite encounter rate, shifting a given sex towards the right in Figure 3. The optimal investment changes accordingly. While increasing *α* lowers optimal investment, the effect of increased parasite encounter rate is system-specific, as explained in previous findings (Figure 1-2). Consequently, what matters for the final prediction is the location of a sex in the parameter space (Figure 3, arrows). If the (more strongly) sexually selected sex encounters parasites more often, it may invest either more (orange arrow) or less (blue arrow) in immunocompetence than the opposite sex. Thus, males, despite facing stronger sexual selection, may invest more in parasite defence, if the system favours improved defences due to more frequent encounters. However, when futility applies, both aspects of sexual selection (its strength per se, and increase in parasite encounter rates due to sexually selected traits) are expected to select for weaker immunocompetence in males than in females (green arrows).

## Discussion

The concept of “futility” explains a counterintuitive phenomenon, where a host neglects immunocompetence when intuitively it would be most needed. Typically, the axis in question is parasite abundance: futility means a negative relationship between (some values of) abundance and evolved resistance (Walsman *et al*., 2023). Broadly speaking, a “resistance-is-futile” strategy applies to ineffective defence mechanisms, predicting weaker or possibly abandoned investments into defence (Jokela *et al*., 2000). In our study, we showed that resistance and tolerance differ in the conditions that create futility.

In addition to resistance diminishing with very high parasite abundance, our study identifies another axis of futility: with tolerance as the defence trait, parasite virulence — not abundance — serves as the axis of futility. Interestingly, ours is not the first to show futility as a response to parasite traits beyond abundance. Best *et al*. (2010) modelled a special case of castrating parasites, distinguishing between two traits termed ‘resistance’ and ‘tolerance’, but with important terminological differences between their study and ours. Their resistance does not prevent infection, instead it is assumed to lower the parasite growth rate, influencing virulence. Thus, once the terminology is accounted for, their finding of “futile resistance” aligns well with virulence-tolerance futility from our model.

Intriguingly, if we assume that immunocompetence aids both resistance and tolerance, neither axis of futility remains; instead, a host will intensively fight a severe parasite burden (abundant, virulent parasites).

How realistic is our model? A potential limitation of our study is that our models considered immunocompetence as a single trait (albeit sometimes with pleiotropic effects), while in reality resistance and tolerance may compete for resources within an organism, as inferred from physiological mechanisms in birds (Arriero *et al*., 2018), and through experimental challenge of fruit flies with an infectious agent (Vincent & Sharp, 2014).

We assume a trade-off between reproduction and immunity that yields sex-specific optima. There are extreme examples in nature: male semelparity in *Antechinus* is caused by the collapse of male immunocompetence after one intense mating season (Fisher *et al*., 2013). More generally, fighting off infection negatively impacts a host’s reproductive success (as suggested by host traits (Manjerovic & Waterman, 2012), and exposure to infectious agents (Hogg & Hurd, 1995; Fellowes *et al*., 1999, for a review see Schwenke *et al*., 2016). In turn, higher reproductive investment may also suppress immunocompetence (Adamo *et al*., 2001; Ardia *et al*., 2003; Fedorka *et al*., 2004; Uller *et al*., 2020). Our model, where sexual selection decreases optimal investment, is in line with previous studies (Zuk, 1990; Sheldon & Verhulst, 1996; Zuk & Stoehr, 2002).

We add, however, new examples to the set of known theoretical counterexamples, such as the known theoretical finding that if a male needs very good condition to have appreciable mating success, and parasites are a major determinant of condition, males may remain the more immunocompetent sex (Stoehr & Kokko, 2006). In our case, sex difference in optimal immunity levels occur when sexes differ in the strength of sexual selection and/or the parasite encounter rate. If parasites are not abundant enough for futility to emerge, the more sexually selected sex is predicted to have lower immunocompetence when sexual selection is the sole difference between sexes; but this sex can switch to higher immunocompetence if sexual selection comes with behaviours increasing parasite encounter rates. In “futile” scenarious, the sexually selected sex is much more clearly predicted to have lower immunocompetence, regardless of parasite encounter rate.

A higher parasite burden in males (Poulin, 1996) may stem either from sex-specific parasite behaviour (Duneau *et al*., 2012; Duneau & Ebert, 2012) or from males offering an easier habitat for the parasites, reflecting sexually divergent immunocompetence (Bacelar *et al*., 2011; Foo *et al*., 2017). Documenting parasite encounter rates is probably easier than sex-specific virulence (Hall & Mideo, 2018). Note, however, that while we might expect higher parasite encounter rates for males (e.g., with male-biased mate-searching, Li & Kokko, 2019), females face unique challenges too, e.g. pregnancy-associated transmission risk (Mitchell *et al*., 2022), traumatic insemination (Reinhardt *et al*., 2003), or diet differences (De Lisle & Rowe, 2021).

To keep our focus on futility, we did not consider parasite virulence evolution (Cousineau & Alizon, 2014; Gipson & Hall, 2016), or a specific form of sexual selection, where favourable sexually selected traits correlate with superior resistance genes (Taskinen & Kortet, 2002; Dunn *et al*., 2013). Stoehr & Kokko (2006) considered immunity investment’s potential to increase mating success, yet without an explicit focus on females preferring resistant males. If females mate preferably with males resisting infection, or tolerant males simply persist longer in the mating pool (an idea akin to ‘stamina’, Kovalov & Kokko, 2022), dynamics are likely to become more complicated; parental care (Venkateswaran *et al*., 2021; Wittman & Cox, 2021) and sexually transmitted disease (Keiser *et al*., 2020) offer further challenges.

Overall, we have shown an interesting non-additivity: two axes of futility both disappear if the same immunity trait can help uninfected hosts avoid infection, and improve the performance (here, survival) of infected hosts. Hence, an immune system with pleiotropic effects may avoid scenarios where immune defence becomes futile. Since it is conceivable that mechanisms conferring resistance can also help tolerating the infection once it exists, this may help explain why organisms rarely ‘capitulate’ in the presence of frequently encountered parasites.

## Acknowledgements

V.K. thanks members of the Lüpold’s group at UZH for their valuable comments at different stages of the work on this project.

## Figures & Tables

**Table 1:**
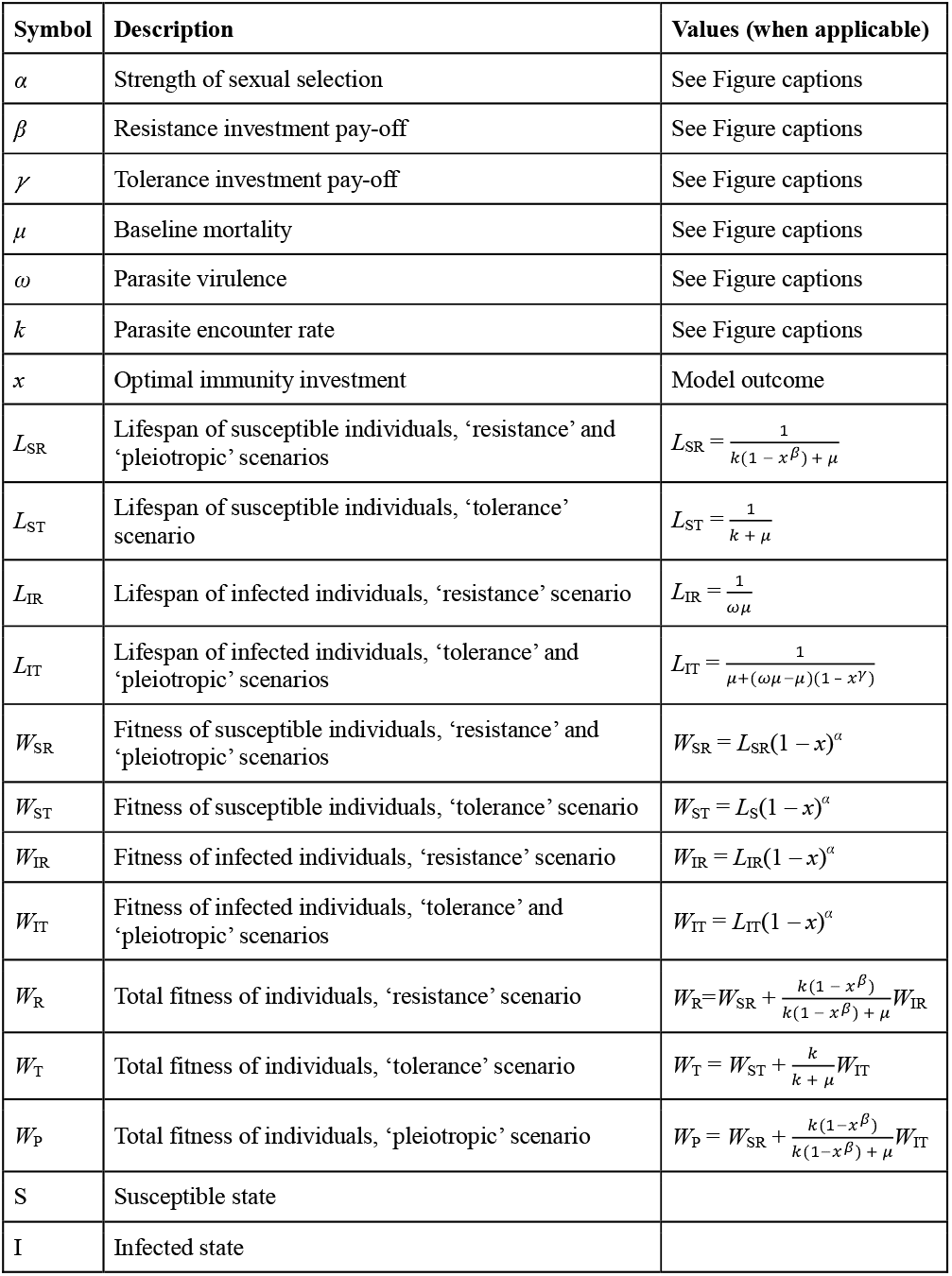
The list of parameters and notions used in the model.

| Symbol | Description | Values (when applicable) |
| --- | --- | --- |
| $\alpha$ | Strength of sexual selection | See Figure captions |
| $\beta$ | Resistance investment pay-off | See Figure captions |
| $\gamma$ | Tolerance investment pay-off | See Figure captions |
| $\mu$ | Baseline mortality | See Figure captions |
| $\omega$ | Parasite virulence | See Figure captions |
| $k$ | Parasite encounter rate | See Figure captions |
| $x$ | Optimal immunity investment | Model outcome |
| $L_{SR}$ | Lifespan of susceptible individuals, ‘resistance’ and ‘pleiotropic’ scenarios | $L_{SR} = \frac{1}{k(1-x^\beta) + \mu}$ |
| $L_{ST}$ | Lifespan of susceptible individuals, ‘tolerance’ scenario | $L_{ST} = \frac{1}{k + \mu}$ |
| $L_{IR}$ | Lifespan of infected individuals, ‘resistance’ scenario | $L_{IR} = \frac{1}{\omega\mu}$ |
| $L_{IT}$ | Lifespan of infected individuals, ‘tolerance’ and ‘pleiotropic’ scenarios | $L_{IT} = \frac{1}{\mu + (\omega\mu - \mu)(1 - x^\gamma)}$ |
| $W_{SR}$ | Fitness of susceptible individuals, ‘resistance’ and ‘pleiotropic’ scenarios | $W_{SR} = L_{SR}(1 - x)^\alpha$ |
| $W_{ST}$ | Fitness of susceptible individuals, ‘tolerance’ scenario | $W_{ST} = L_S(1 - x)^\alpha$ |
| $W_{IR}$ | Fitness of infected individuals, ‘resistance’ scenario | $W_{IR} = L_{IR}(1 - x)^\alpha$ |
| $W_{IT}$ | Fitness of infected individuals, ‘tolerance’ and ‘pleiotropic’ scenarios | $W_{IT} = L_{IT}(1 - x)^\alpha$ |
| $W_R$ | Total fitness of individuals, ‘resistance’ scenario | $W_R = W_{SR} + \frac{k(1-x^\beta)}{k(1-x^\beta) + \mu} W_{IR}$ |
| $W_T$ | Total fitness of individuals, ‘tolerance’ scenario | $W_T = W_{ST} + \frac{k}{k + \mu} W_{IT}$ |
| $W_P$ | Total fitness of individuals, ‘pleiotropic’ scenario | $W_P = W_{SR} + \frac{k(1-x^\beta)}{k(1-x^\beta) + \mu} W_{IT}$ |
| S | Susceptible state |  |
| I | Infected state |  |

