## Supplementary Material for "Sex-specific immunocompetence: resistance and tolerance can both be futile but not under the same circumstances"

* – shared first coauthor

[+] – corresponding author

1 – Department of Evolutionary Biology and Environmental Studies, University of Zurich, Winterthurerstrasse 190, 8057 Zurich, Switzerland

2 – Institute for Organismic and Molecular Evolution and the Institute of Quantitative and Computational Biology, Johannes Gutenberg University of Mainz, Hanns-Dieter-Husch Weg 15, 55128 Mainz, Germany

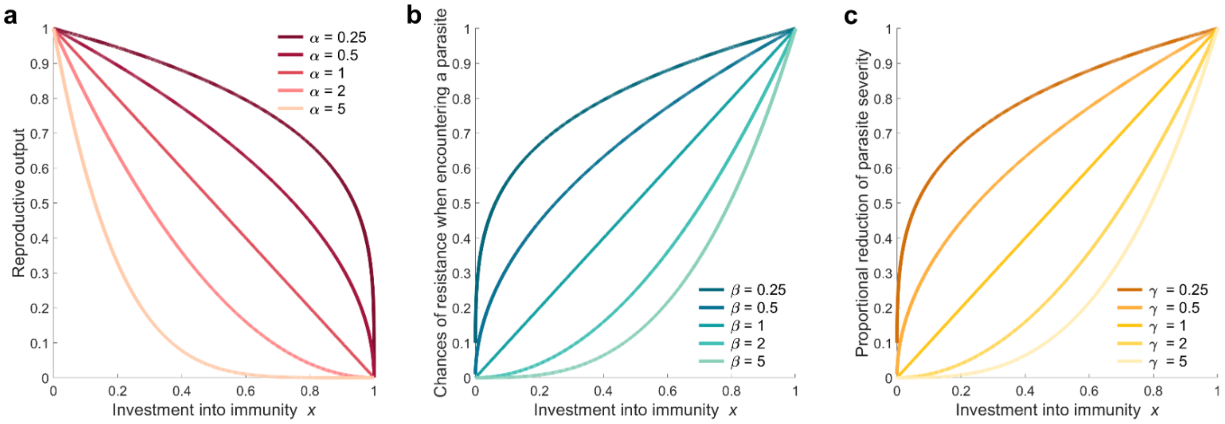


**S1:** *Relationship between investment into immunity and resulting reproductive and immune payoffs.* a) Interaction between immunity investment and resulting reproductive output, depending on $\alpha$. Higher values of $\alpha$ represent a scenario of strong sexual selection. Hosts need to make great energetic investment into reproduction (and in correspondence little investment into immunity) to achieve higher reproductive success. b) Higher investment into immunity corresponds to a greater chance of resisting infection. $\beta$ scales how immunity investment translates into improved resistance. c) Investment into immunity turns into a reduction of parasite severity. The higher the investment, the lower the parasitic virulence a host experiences. $\gamma$ regulates the level to which immunity investment payoff with increased tolerance. With greater $\gamma$ values, investment into immunity need to be high to significantly reduce parasite severity. Note how the immunity mechanisms, scaled by $\beta$ (in b) and $\gamma$ (in c) show the same relationship between investment and payoff for both resistance and tolerance.

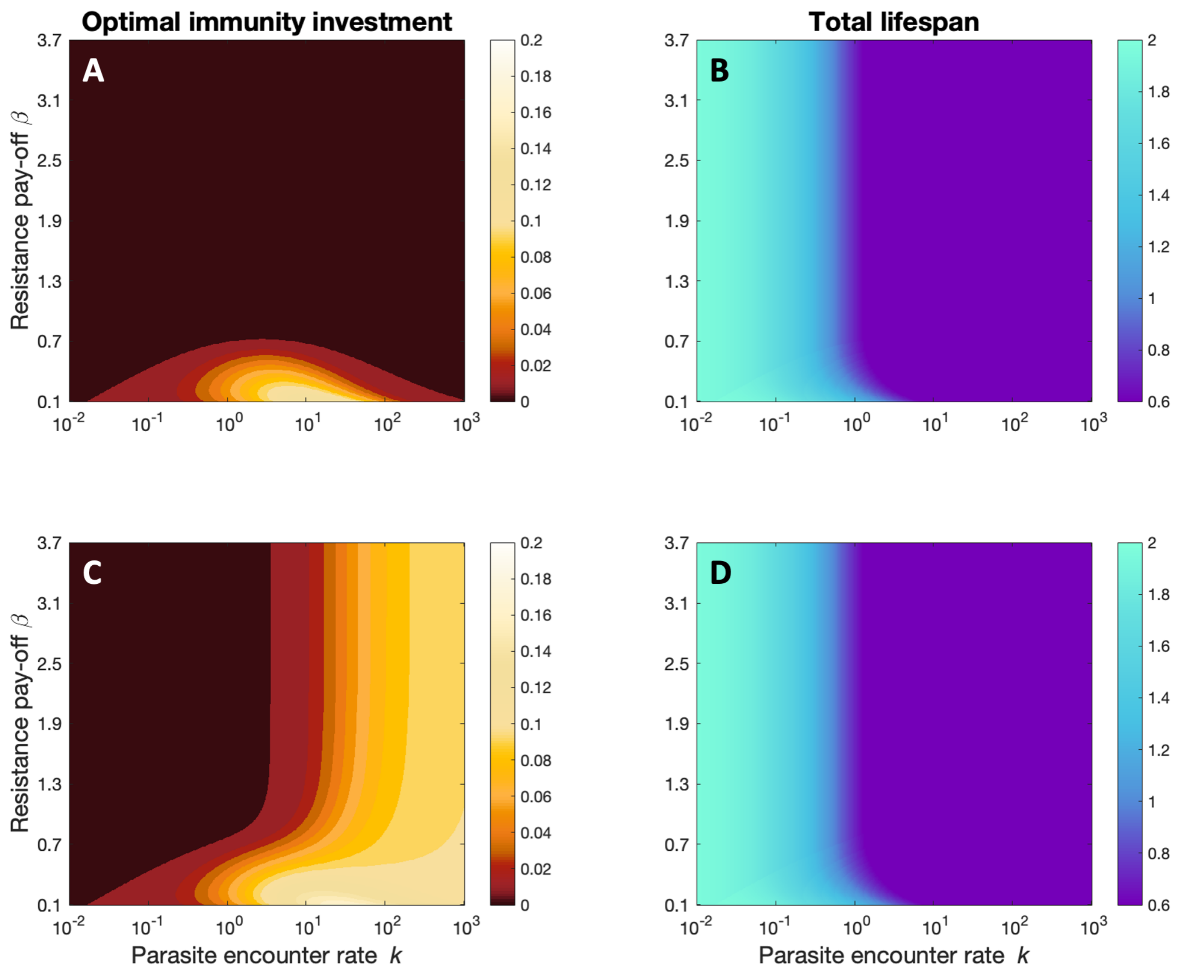


**S2:** *The effect of resistance investment pay-off and parasite encounter rate on optimal immunity investment and total lifespan.* Immunocompetence is assumed to improve resistance (A, B), or both resistance and tolerance (C, D). The investment heatmaps (A, C) show the optimal immunity investment $x$ for any combination of parasite encounter rate $k$ (x-axis, log scaled) and resistance investment pay-off $\beta$ (y-axis), with $x$ represented as a gradient from dark red ($x = 0$) to white ($x \sim 0.99$, the maximum value in this model setup). The lifespan heatmaps (B, D, F) assume individuals invest according to the optimal value $x$, depicting expected lifespan as a gradient from dark violet ($L = 0.6$) to cyan ($L = 2$). Model parameters: $\alpha = 2$; $\beta = 0.1\ldots4$; $\gamma = 0.5$; $k = 10^{-2}\ldots10^{3}$; $\mu = 0.5$; $\omega = 30$.

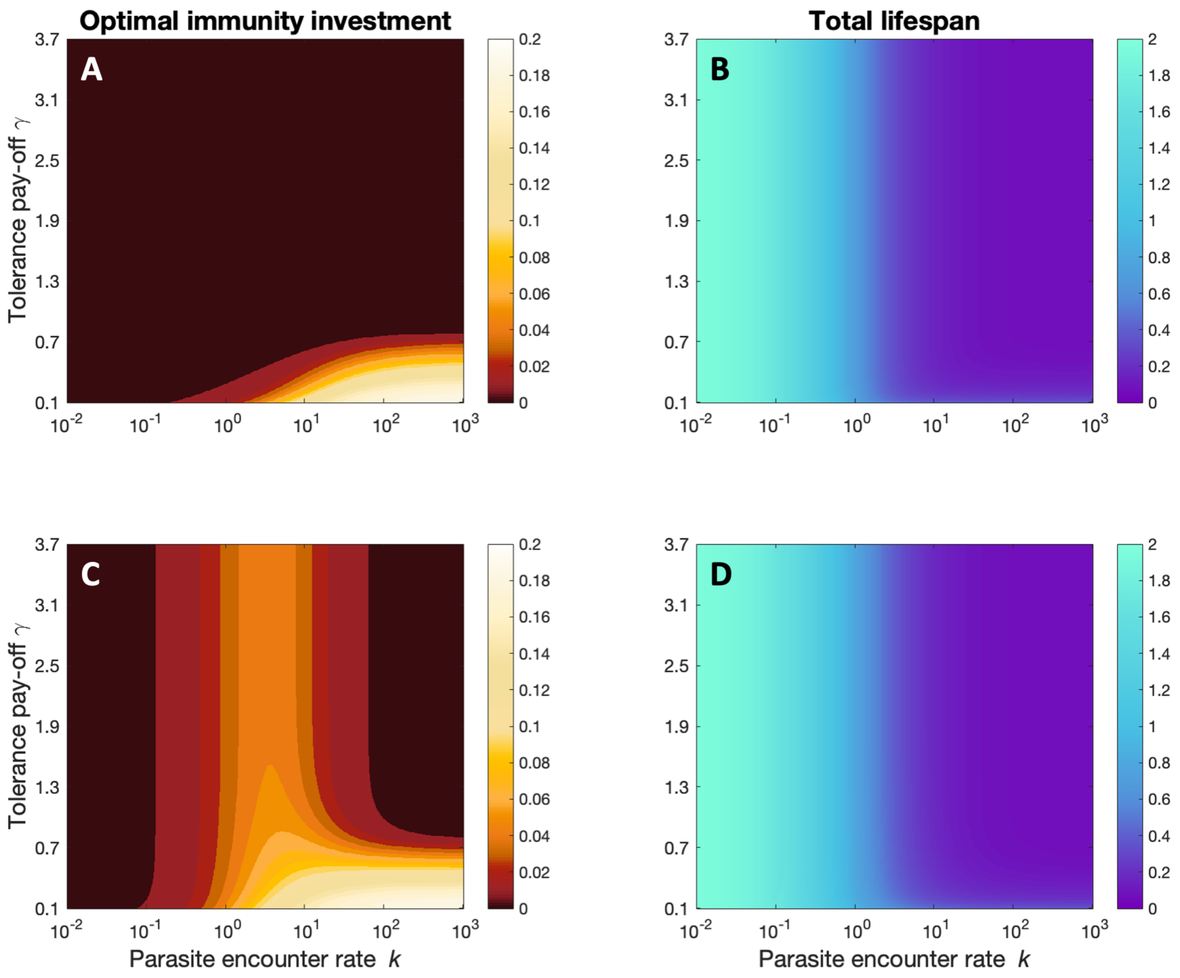


**S3:** *The effect of tolerance investment pay-off and parasite encounter rate on optimal immunity investment and total lifespan.* Immunocompetence is assumed to improve tolerance (A, B), or both resistance and tolerance (C, D). The investment heatmaps (A, C) show the optimal immunity investment $x$ for any combination of parasite encounter rate $k$ (x-axis, log scaled) and resistance investment pay-off $\gamma$ (y-axis) within their value ranges. Immunity investment value $x$ is represented as a gradient from dark red (x = 0) to white (x = 0.2), with $x$ represented as a gradient from dark red ($x = 0$) to white ($x \sim 0.99$, the maximum value in this model setup). The lifespan heatmaps (B, D, F) assume individuals invest according to the optimal value $x$, depicting expected lifespan as a gradient from dark violet ($L = 0.6$) to cyan ($L = 2$). Model parameters: $\alpha = 2$; $\beta = 0.5$; $\gamma = 0.1 \ldots 4$; $k = 10^{-2} \ldots 10^{3}$; $\mu = 0.5$; $\omega = 30$.

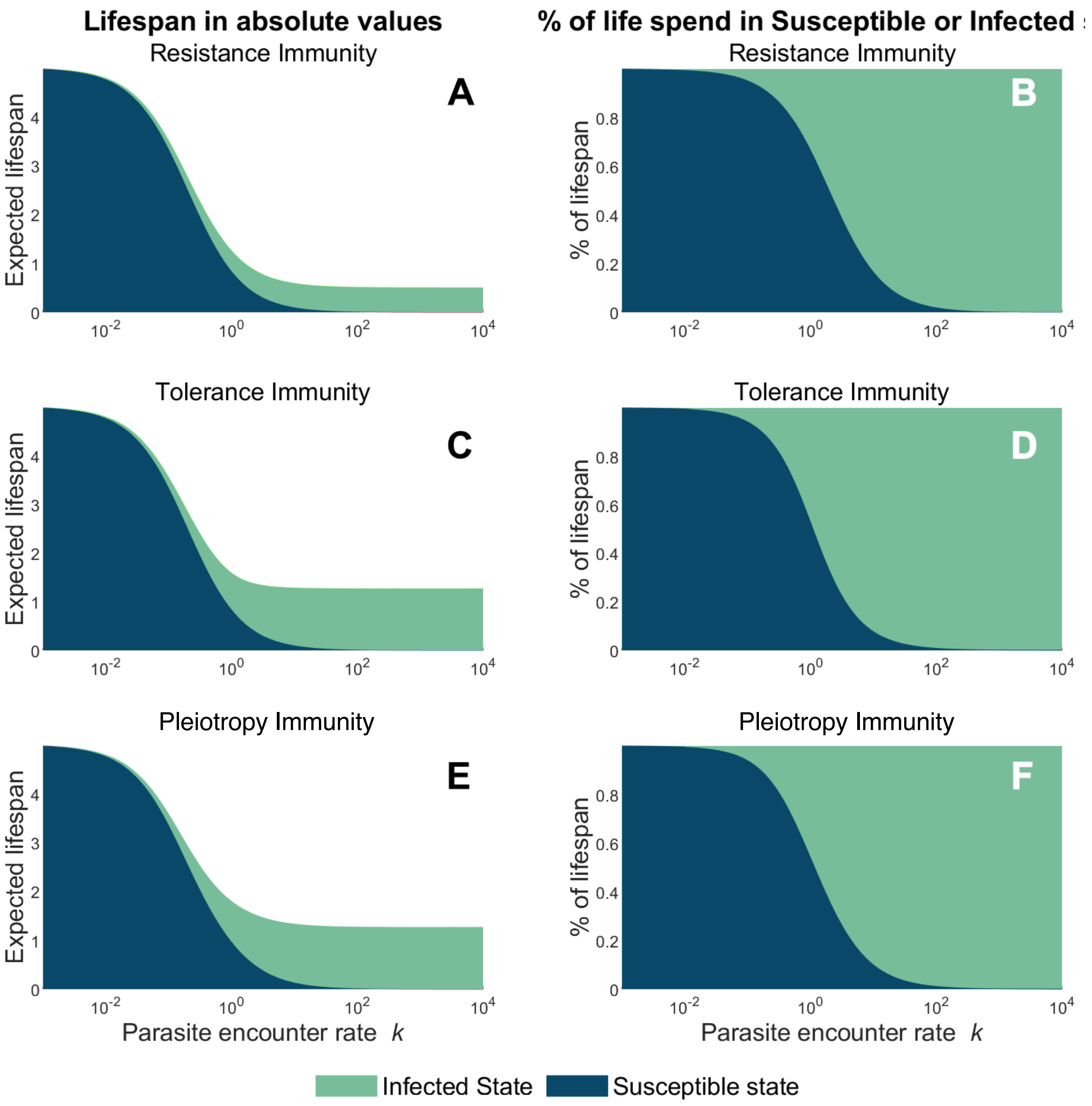


**S4:** *Expected Lifespan resulting from optimal immunity investment depending on parasite abundance.* Immunocompetence is assumed to improve resistance (A, B), tolerance (C, D), or both (E, F). Green-colored areas correspond to time an organism is expected to spend in the infected state, while blue-colored areas mark time spent in the susceptible state. Expected total lifespan changes with parasite encounter rate $k$ in absolute values (A, C, E). The percentage of lifetime spent in the susceptible or infected state similarly changes with increasing parasite encounter rate $k$ (B, D, F). Model parameters $\alpha$=1.5, $\beta$=0.8, $\gamma$=0.25, $\Omega$=10 and $\mu$=0.2.
